# Implicit temporal predictability enhances pitch discrimination sensitivity and biases the phase of delta oscillations in auditory cortex

**DOI:** 10.1101/410274

**Authors:** Sophie K Herbst, Jonas Obleser

## Abstract

Can human listeners use implicit temporal contingencies in auditory input to form temporal predictions, and if so, how are these predictions represented endogenously? To assess this question, we implicitly manipulated temporal predictability in an auditory pitch discrimination task: unbeknownst to participants, the pitch of the standard tone could either be deterministically predictive of the temporal onset of the target tone, or convey no predictive information. Predictive and non-predictive conditions were presented interleaved in one stream, and separated by variable inter-stimulus intervals such that there was no dominant stimulus rhythm throughout. Even though participants were unaware of the implicit temporal contingencies, pitch discrimination sensitivity (the slope of the psychometric function) increased when the onset of the target tone was predictable in time (N = 49, 28 female, 21 male). Concurrently recorded EEG data (N = 24) revealed that standard tones that conveyed temporal predictions evoked a more negative N1 component than non-predictive standards. We observed no significant differences in oscillatory power or phase coherence between conditions during the foreperiod. Importantly, the phase angle of delta oscillations (1–3 Hz) in auditory areas in the post-standard and pre-target time windows predicted behavioral pitch discrimination sensitivity. This suggests that temporal predictions are encoded in delta oscillatory phase during the foreperiod interval. In sum, we show that auditory perception benefits from implicit temporal contingencies, and provide evidence for a role of slow neural oscillations in the endogenous representation of temporal predictions, in absence of exogenously driven entrainment to rhythmic input.

## Introduction

The human brain is constantly forming predictions about its environment (Friston, 2005; Rao and Ballard, 1999), which concern the *where* and *what* of future events, but also the *when* (Arnal and Giraud, 2012; Coull and Nobre, 1998; Nobre et al., 2007; Nobre and van Ede, 2018; Rimmele et al., 2018). To predict when future events will occur, temporal statistics of sensory input are extracted and translated into temporal predictions that benefit perception and action. Yet little is known about how endogenous temporal predictions are formed from temporal regularities in sensory input, and how they are represented in human brain dynamics.

Temporal predictions are often enabled by an isochronous periodic structure of sensory inputs, to which we will refer as *rhythmic* temporal predictions in the following. Rhythmic input structure has been shown to improve detection performance and speed (Henry and Obleser, 2012; Lawrance et al., 2014; Rimmele et al., 2011; Stefanics et al., 2010; Wright and Fitzgerald, 2004). Fewer studies have shown that rhythmic temporal predictions can also improve perceptual sensitivity (i.e. discrimination performance) in the auditory (Jones et al., 2002; Morillon et al., 2016; Schmidt-Kassow et al., 2009; but see Bauer et al., 2015), as well as the visual domain (Cravo et al., 2013; Rohenkohl et al., 2012). It is, however, not trivial to disentangle mechanistic input-driven alignment of neural activity to rhythmic input from an internalized and endogenously activated representation of temporal predictions (Haegens and Golumbic, 2017; Rimmele et al., 2018; van Wassenhove, 2016).

To assess the endogenous representation of temporal predictions, devoid from the representation of the periodic structure of sensory input, we here induced temporal predictability by manipulating the temporal statistics in a so-called foreperiod paradigm (Niemi and Näätänen, 1981; Woodrow, 1914). This type of manipulation has been shown to increase visual perceptual sensitivity (Correa et al., 2004, 2005; Cravo et al., 2011; Rolke and Hofmann, 2007). In audition, temporally predictable foreperiods have been found to speed up stimulus processing (Bausenhart et al., 2007) and improve short-term memory performance (Wilsch et al., 2018, 2014). Morillion et al. (2016) reported an increase in auditory sensitivity, inducing aperiodic but ordered temporal regularities.

Importantly, forming temporal predictions does not require conscious awareness of the temporal structure, but can occur implicitly (Cravo et al., 2011; Herbst and Obleser, 2017). While some previous studies used explicit temporal prediction tasks, in which temporal regularities were fully disclosed to participants (Stefanics et al., 2010), here we aim at studying the automatic extraction of temporal predictions from sensory environments, to mimic naturalistic settings.

To assess an endogenous representation of temporal predictions, we investigated the hypothesis that slow neural oscillations (in the delta/1–3 Hz and theta/4–7 Hz frequency bands) implement temporal predictions via endogenous phase-resetting and -shifting mechanisms. This hypothesis can be drawn back to the influential proposal of *Dynamic Attending in Time* (DAT; Jones, 1976; Large and Jones, 1999), suggesting that (auditory) attention fluctuates in phase with rhythmic input. A neural implementation of dynamic attending has been postulated through phase-locking of neural delta oscillations to rhythmic inputs, also termed entrainment. Entrainment reflects an internalization of the exogenous temporal structure, to align the most efficient brain states for sensory processing to the most likely time points for stimulus occurrence (Lakatos et al., 2008; Schroeder and Lakatos, 2009). Behaviourally, this results in fluctuations of performance in phase with the oscillation (Barczak et al., 2018; Besle et al., 2011; Cravo et al., 2013; Kösem et al., 2014; Lakatos et al., 2008; Morillon and Baillet, 2017; Schroeder and Lakatos, 2009; Stefanics et al., 2010).

It is currently an open question to what extend entrained delta oscillations are a generic signature of processing rhythmic input, versus specifically represent a neural implementation of temporal predictions. Important evidence for a specific role of endogenous delta oscillations for temporal processing in audition comes from two studies showing that auditory processing fluctuates with the phase of spontaneously present delta activity in auditory cortex, in absence of rhythmic stimulation (Henry et al., 2016; Kayser et al., 2015). Furthermore, previous studies have shown that entrainment is subject to top-down modulation, as phase coherence of slow oscillations in anticipation of temporally predictive input scales with the strength of temporal predictions (Breska and Deouell, 2017; Cravo et al., 2013; Stefanics et al., 2010).

As a means to experimentally separate endogenous delta oscillations from exogenous stimulus rhythms and the resulting entrainment of neural oscillations, studies have started to test whether the phase of an ongoing oscillation can be aligned in a top-down manner to an expected point in time, without an entraining stimulus structure (Cravo et al., 2011; Herbst and Obleser, 2017; Solís-Vivanco et al., 2018). To our knowledge, only one study in the visual domain reported an effect of increased phase coherence in single-interval temporal predictions (Cravo et al., 2011, theta band). Furthermore, a recent study (Barne et al., 2017) showed that delta phase in the target-onset time window reflects adjustments to previously encountered violations of temporal predictions in an explicit timing task.

Here, to investigate the role of slow oscillatory dynamics for an endogenous representation of temporal predictions in auditory inputs, in absence of a rhythmic structure, we implicitly associated temporal predictability to a sensory feature of the standard tone in an auditory pitch discrimination task: the standard’s pitch could be deterministically predictive of the onset time (but not the pitch) of the target tone, or convey no predictive information. Temporally predictive and non-predictive conditions were presented interleaved in one stream, and separated by variable inter-stimulus intervals such that there was no dominant stimulus rhythm throughout.

We hypothesized that, behaviourally, temporal predictability would increase pitch discrimination sensitivity, assessed via the slope of the psychometric function. In the concurrently recorded EEG data, we expected to see indices of temporally predictive processing in the auditory evoked potential, namely the N1 and P2 components. Based on the current literature the expected direction of the effect is not clear (Lange, 2013). Furthermore, we expected to confirm a hypothesized role of delta oscillations in temporal prediction, surfacing as enhanced phase coherence in the temporally predictive condition (Stefanics et al., 2010), or a direct relationship between delta phase and our behavioral measures (Cravo et al., 2013).

## Methods

### Participants

In total, 51 participants were tested (23.6 years on average (SD = 3.5), 28 female, 6 left handed), 26 of which also underwent electroencephalography (EEG). All participants signed informed consent and received either course credit or payment for their participation (8 € per hour). The study was approved by the local ethics commit tee at the University of Lübeck (17-154). We excluded two of the participants who only underwent the behavioral testing, because of ceiling effects (their slopes for the psychometric function in one of the two conditions exceeded the mean of the slope distributions of all participants by more than 2.5 standard deviations). Furthermore, we excluded the EEG data from two participants who had blinked in synchrony with the auditory stimulation and for whom we were not able to separate blinks from the auditory evoked potentials during EEG preprocessing. The behavioural data of these two participants were kept in the analyses.

### Stimuli and Procedure

The experiment was conducted in an electrically shielded sound-attenuated EEG booth. Stimulus presentation and collection of behavioural responses was achieved using the Psychophysics Toolbox (Brainard, 1997; Pelli, 1997) under Windows 7. Responses were collected on a standard keyboard. All participants were instructed to use the index and middle fingers of the right hand.

Participants performed a pitch discrimination task, comparing tone pairs embedded in noise, as illustrated in Figure 1A. They were instructed to indicate after each tone pair whether the second tone was lower or higher than the first.

**Fig 1.**
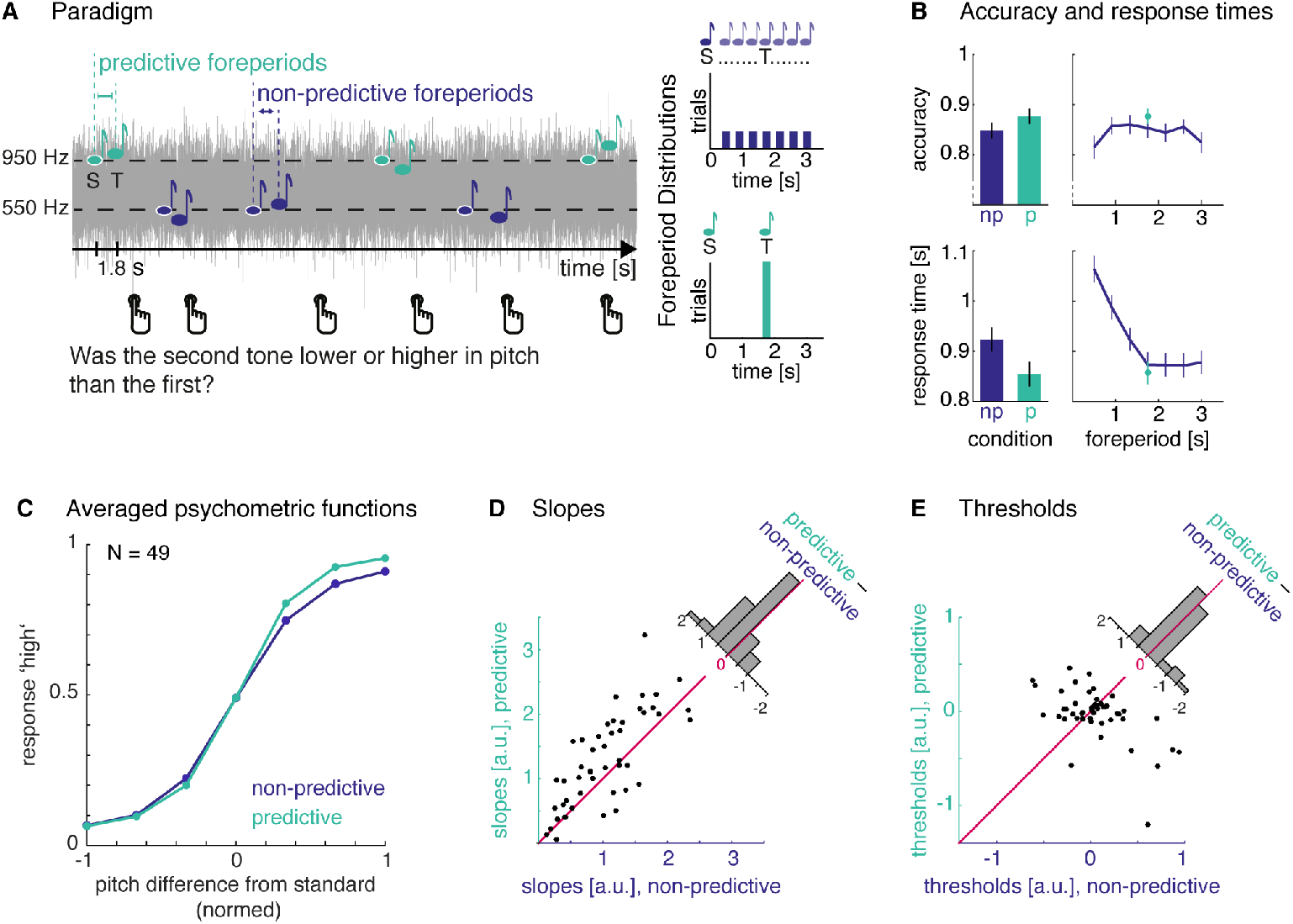
Paradigm and Behavioural Results. **A. Paradigm**: Tone-pairs were presented embedded in low-pass filtered white noise. Participants’ task was to judge whether the target tone (T) was lower or higher in pitch than the preceding standard (S). Unbeknownst to participants, the pitch of the standard tone was associated with predictive (green) or non-predictive foreperiod intervals (blue). For the non-predictive condition, foreperiods were drawn from a uniform distribution (upper right panel), while for the predictive condition, foreperiods were fixed at 1.75 s (lower right panel). **B. Accuracy and response times**: Top: Accuracy improved significantly in the predictive condition (left panel), which was nominally also true at the intermediate foreperiod only (right panel). Bottom: Response times were faster in the predictive condition (left panel). The difference was driven by slower response times at short foreperiods in the non-predictive condition (right panel) **C. Averaged psychometric functions**: The slope of the psychometric function was steeper in the predictive compared to the non-predictive condition. There were no differences in threshold, nor the lower or upper asymptotes. **D. Slopes for single participants**: for the non-predictive (x-axis) versus predictive (y-axis) conditions. **E. Thresholds for single participants**: for the non-predictive (x-axis) versus predictive (y-axis) conditions. a.u. stands for absolute units.

A black fixation cross was displayed on gray background throughout the whole block. Auditory stimuli were delivered via headphones (Sennheiser HD 25-SP II). Lowpass (5kHz) filtered white noise was presented constantly throughout each block, at 50 dB above the individual sensation level, which was determined for the noise alone at the beginning of the experiment using the method of limits. Pure tones of varying frequencies (duration 50 ms with a 10 ms on- and offset ramp), were presented with a tone-to-noise ratio fixed at −18 dB relative to the noise level.

The first tone, to which we will refer as the *standard* in the following was always at one of two frequencies: 550 or 950 Hz. The second tone, the *target,* was varied in individually predetermined steps around its respective standard. The same step size was used for both standards, but logarithmically transformed and multiplied with the standard frequency, to obtain a log-spaced frequency scale around each standard. To predetermine the step size, each participant was first presented with one experimental block, containing all tone steps to familiarize themselves with the task. Then, a second block was performed, and if pitch discrimination performance across steps was below 65%, the tone-steps were increased, which was repeated up to three times. All participants reached the minimum performance level after minimally two and maximally four rounds of training. As a result of this procedure, the average lowest target tone presented with the 550 Hz standard was 508.3 Hz (range 490.0–519.1 Hz), and the highest target tone 595.3 Hz (range 582.7–617.4 Hz); the lowest target tone presented with the 950 Hz standard was 878.0 Hz (range 846.4–896.7 Hz), and the highest target tone 1028.3 Hz (range 1006.5–1066.3 Hz). The high and low tones never overlapped. In the behavioural experiment, eleven tone frequencies were used from the lowest to highest tone, including the standard; in the EEG experiment we used 7 discrete frequencies.

Critically, and unbeknownst to participants, we manipulated the interval between standard and target tones, the *foreperiod,* by either pseudo-randomly drawing foreperiods from a discretized uniform foreperiod duration (11 foreperiods in the behavioral experiment and 7 in the EEG experiment, all ranging from 0.5–3 s, blue distribution in Figure 1 A), or used the same foreperiod duration (1.75 s, green distribution in Figure 1 A). This resulted in one condition in which the target onset was perfectly predictable in time, the *predictive condition,* and one condition in which the target onset was maximally jittered, the *non-predictive condition.* To allow participants to implicitly dissociate the conditions, the foreperiod distributions were associated with one of the standard pitches, for example for one participant the 550 Hz standard was always followed by a predictive foreperiod and the 950 Hz standard was always followed by a non-predictive foreperiod. The assignment was counterbalanced over participants. The two conditions were presented interleaved, such that participants had to encode the standard pitch on each trial. Importantly, the manipulation of foreperiod intervals was strictly implicit, and participants were not informed about it.

To avoid build-up of a rhythm over trials, the inter-stimulus interval between a target tone and the standard tone of the next trial was drawn from a truncated exponential distribution (mean 1.5 s, truncated at 3 s) added to a minimum interval of 3 s (resulting in values between 3–6 s). After the target tone, participants had 2 s to respond. The stimulation continued automatically, even if no response was given.

One block consisted of 22 trials in the behavioural (one repetition per tone step and condition), and 56 trials in the EEG experiment (4 repetitions per tone step and condition). In the behavioural experiment participants performed 20 blocks (440 trials), and in the EEG experiment minimally 12 and maximally 15 blocks (672–840 trials). Between blocks, participants could take breaks of self-determined length. Feedback was given per trial during the training, and at the end of each block (as proportion of correctly answered trials) during the main experiment.

After the experiment, all participants were asked the same four questions by the experimenter. First, the experimenter asked whether participants had noticed that the interval between the first and second tone of a pair was variable. Second, they were asked to describe whether they noticed any systematic variation therein. Third, they were told that either the low or high tones were always presented with the same separating interval and asked whether they noticed this. Fourth, they were asked to guess whether in their case the low or high pitch tones were the ones presented with the constant interval. Finally, they filled in a musicality survey (Schaal et al., 2014). The full experimental session lasted about 2.5 h.

### EEG recording and preprocessing

EEG was recorded with 64 electrodes Acticap (Easy Cap) connected to an ActiChamp (Brain Products) amplifier. EEG signals were recorded with the software Brain Recorder (Brain Products) at a sampling rate of 1 kHz, using no online high-pass filter and a 200 Hz low-pass filter. Impedances were kept below 10 kΩ. Electrode TP9 (left mastoid) served as reference during recording. Electrode positions were digitized.

EEG data were analysed using the Fieldtrip software package for Matlab (MATLAB 2016a, MATLAB 2017a), and the lme4 package in R (Bates et al., 2015; R Core Team, 2016). First, we re-referenced the data to linked mastoids. Then we applied a low-pass filter to the continuous data (cut-off 45 Hz, two-pass, transition bandwidth 3 Hz; firws filter from the firfilt plugin, Widmann et al., 2015). No high-pass filter was applied. For the time-frequency analysis, we produced a parallel version of the data, that was not filtered during pre-processing. Filtering two-pass as done for the analyses of event-related potentials might smear data back in time, which would be problematic for analyses in the pre-target time window (Rousselet, 2012; Zoefel and Heil, 2013). Filtering the data only in the forward direction, however, leads to phase shifts (Widmann et al., 2015) which we wanted to avoid for the phase angle analyses.

Next, we epoched the data around the standard tone onset (–3 to 6 s), and downsampled to 100 Hz. All data were visually inspected to mark bad channels that were interpolated (1.2 channels per participant on average). Then ICA were computed using the ‘runica’ algorithm, with the number of output components adjusted by subtracting the number of bad channels. Blinks, muscular artefacts, and unspecific noise occurring temporarily in a channel or trial were excluded, using the semi-automatic inspection of ICA components provided by the SASICA toolbox for fieldtrip (Chaumon et al., 2015) and removal of these (on average 33.7 components per participant).

## Analyses

### Analyses of the behavioural data

We analysed accuracy as proportion correct (after removing trials in which the standard and target were equal in pitch) and response times, defined as the interval between the onset of the target tone and the registered button press. Response times shorter than 0.2 s were considered outliers and removed. We compared accuracy and response times between conditions and over foreperiods for the non-predictive condition. Tone-steps and foreperiods used in the behavioral experiment were binned to reduce the 11 steps to 7 to match the steps in the EEG-experiment, by averaging the second and third, fourth and fifth, as well as the seventh and eight and ninth and tenth tone steps.

To obtain a measure of pitch discrimination sensitivity, we fitted psychometric functions to model participants’ responses in the pitch discrimination task, using bayesian inference, implemented in the *Psignifit toolbox* for Matlab (Version 4, Schütt et al., 2016). The psychometric function describes the relationship between the stimulus level (on the abscissa, here: the difference in pitch between the target and the respective standard tone) and the participant’s answer (on the ordinate, here: proportion of trials on which the target pitch was judged as higher).To accommodate the different standard tones per condition, and the individual pitch steps obtained during the training, we transformed the discrete tone frequencies per participant and condition to 11, or respectively 7 linearly spaced steps from −1 to 1, with −1 and 1 reflecting each participant’s extremest tones, and 0 being the pitch of the standard tone.

To select the options for the psychometric function (logistic versus cumulative normal function, number of free parameters), we assessed deviance pooled for both conditions. Deviance reflects a monotonic transformation of the log-likelihood-ratio between the fitted model and the saturated model (a model with no residual error), allowing for an absolute interpretation, or a comparison between different models (Wichmann and Hill, 2001). The best fits (i.e. lowest deviance, 3.80 for the best model) were obtained by fitting a cumulative normal function with four free parameters: threshold, slope, lower, and upper asymptote.

For a yes-no-task as the one used here, the threshold parameter indicates the stimulus level at which a participant is as likely to judge the stimulus as ‘low’ or ‘high’. Divergence from the actual midpoint of all stimulus levels (here: 0) can thus be interpreted as a response bias. The slope parameter reflects the amount of stimulus change needed to distinguish between low and high tones, and can be interpreted as the sensitivity of the listener. The lower asymptote indicates the proportion of answering ‘high’ for the lowest pitches in the tested range, and the upper asymptote the proportion of answering ‘low’ for the highest pitches, that is they reflect the errors made by the listener at different target tone frequencies.

We used Psignifit’s default priors for the threshold, slope, guess, and lapse-rates, based on the given stimulus range (Schütt et al., 2016, p.109). Psignifit’s version 4 fits a beta-binomial model (instead of a binomial model), which assumes that the probability for a given proportion of answers is itself a random variable, drawn from a beta distribution. This has been shown to provide better fits for overdispersed data, that is data in which answer probabilities over blocks and trials are not independent as assumed by the conventional model.

We fitted psychometric functions to each individual’s data separately per condition and compared the resulting parameters between conditions (threshold, slope, guess- and lapse rates) using two-sided t-tests. Additionally, we calculated Bayes Factors for all statistical tests, using the *Bayes Factors* package for Matlab (Rouder et al., 2009).

Additionally, we computed a logistic regression on the single-trial responses of the pitch-discrimination task, to parallel the analysis of delta phase angles performed for the EEG (see below). Pitch difference and condition were used as interacting fixed effects (with random intercepts and random slopes for both predictors and their interaction), using the lme4 package in R (function *glmer,* Bates et al., 2015) with a binomial link function.

### Event-related potentials

We examined the time-domain data with respect to responses evoked by standard and target tones, contrasting the predictive and non-predictive condition. For the standard-evoked response, we detrended the data based on the whole epoch and applied baseline correction from −0.1 to 0 s pre-standard. We only examined the time-window between standard onset and 0.5 s after, because this was the maximal interval in which no target events occurred (earliest target onset was 0.5 s in the non-predictive condition). For the target-evoked response, we first applied detrending and the same pre-standard baseline to standard-locked epochs, and then re-epoched to the target event. We examined the time interval from −0.5 to 0.5 s around the target event. We averaged over trials within participants and condition, and then over participants, to obtain the average event-related potential (ERP, depicted in Figure 2).

**Fig 2.**
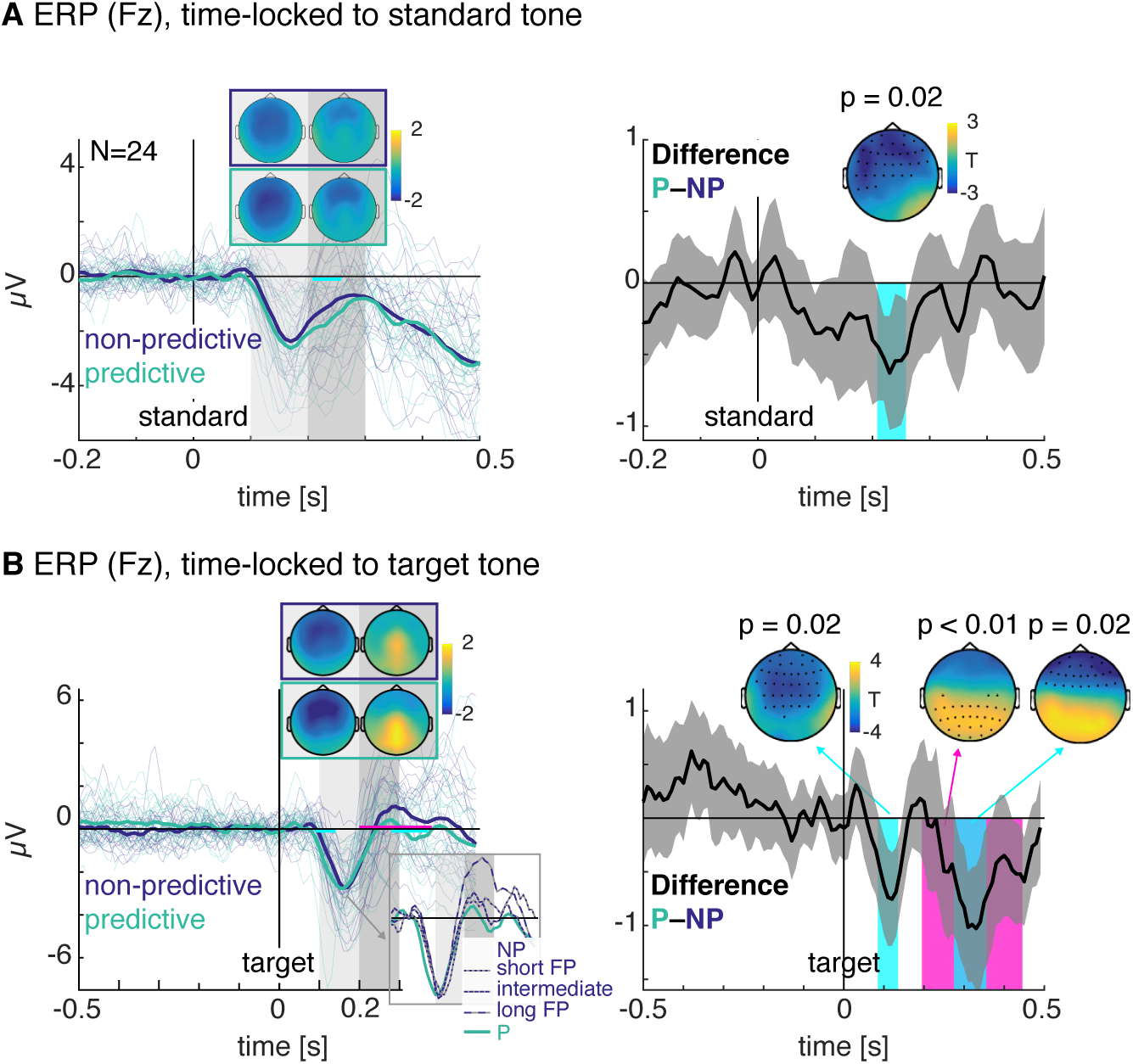
Event-related potentials (ERP). **A. ERPs time-locked to the standard tone**: Left: The predictive condition (green line) evoked a more negative N1 than the non-predictive condition (blue line). The fine blue and green lines depict single participants’ ERPs. The inset shows the topographies in the time windows of 0.1–0.2 s and 0.2–0.3 s for both conditions separately. Right: condition difference. The grey shades indicates the two-sided 95% confidence interval, estimated from the t-distribution. The cyan shade marks the time points at which a significant condition difference occurred, and the topography shows the scalp distribution of the activity during these time windows. Channels at which the difference was significant are marked in black. **B. ERPs time-locked to the target tone**: Left: The predictive condition (green line) evoked an earlier N1 than the non-predictive condition (blue line). The upper inset shows the topographies in the time windows of 0.1–0.2 s and 0.2–0.3 s for both conditions separately. The lower inset exemplary depicts the target-evoked ERP for the 20% longest, intermediate, and 20% shortest foreperiods. Right: condition difference. The cyan and pink shades mark the time points at which a significant condition difference occurred, and the topographies show the scalp distributions of the activity during these time windows.

To test for statistically significant differences in the time-domain data, we applied cluster permutation tests on two levels. First, we contrasted trials from the non-predictive and predictive condition within each participant using independent samples regression implemented in FieldTrip (ft_timelockstatistics). This resulted in regression coefficients (betas) for each time-electrode data point for the ERPs. Next, the group-level analysis was performed with a dependent samples t-test to contrast the betas from the subject-level analysis against zero. A permutation test (5000 Monte Carlo random iterations, minimum of three neighbouring channels to count as a cluster) was performed with cluster-based control of type I error at a level of α=0.05 as implemented in FieldTrip. The condition assignment (i.e. whether the predictive condition was presented at the low or high pitch tones) was added as a control variable. This analysis resulted in time-electrode clusters exhibiting significant condition differences in the ERPs.

### Time-frequency representations

Time-frequency representations were computed for epochs time-locked to the standard tones, separately for the predictive and non-predictive condition. We performed this analysis on trials with foreperiods equal or longer than 1.75 s only to avoid evoked activity from target onsets occurring early in the non-predictive condition. We matched the smaller number of trials available from the non-predictive condition, by randomly sampling the same number of trials from the predictive condition. To obtain stable results, we repeated the random sampling 50 times and averaged over the resulting time-frequency representations. Additionally, we ruled out potential back-smearing of evoked activity related to target-onset by replacing all data points after 1.75 s by the value at this time point for the respective trial and channel before performing the time-frequency transformation.

Data were transformed to time-frequency representations for frequencies ranging from 0.5 to 34.5 Hz (linear steps, 1 Hz) and time points between −0.5 to 2.5 s, using convolution with a single adaptive Hanning taper with frequency-dependent time windows (increasing linearly from 2 to 4 cycles per frequency). To provide sufficiently long data epochs for the lowest frequencies, we appended the epochs (–3 to 6 s, time locked to the standard tone) with their inverted and right-left flipped version to the left and right before applying the time-frequency transform.

Power estimates were extracted as the squared modulus of the complex-valued Fourier spectra and baseline corrected to relative change (first subtracting, then dividing by the trial-average baseline value per frequency) using the condition average in the interval from −0.5 s to standard onset. Inter-trial phase coherence (ITC) was extracted as the magnitude of the amplitude-normalized complex values, averaged across trials for each time-frequency bin and channel.

Statistics were performed in the time-window between 0 to 1.7 s post standard onset and for all frequencies jointly. For power, we used a two-level procedure as described for the ERPs (but using ft_freqstatistics, 1000 permutations). For the ITC, we only computed statistics on the second-level condition differences since it represents a measure that already combines single trials. An additional, hypotheses-driven cluster test for power and ITC effects was performed, restricted to the delta band (1 to 3 Hz).

### Delta phase angle analyses

A timing mechanism that predicts the onset of the target tone would have to be initiated by the standard tone which serves as a temporal cue. Therefore, we examined the data for any signatures of such a mechanism in the phase of the delta band (see Figure 4B for a schematic depiction). To not confound target evoked activity with pre-target activity, we used the same version of the data as for the time-frequency transformations described above, to which no filters had been applied during preprocessing. Target-onset ERPs were muted (as described above) from the time point of target onset on each trial (1.75 s in the predictive condition and 0.5 to 3 s in the non-predictive condition). To reduce the dimensionality of the data, and to focus our analysis on auditory activity, we computed a weighted average of single electrodes at each time point. The weights reflected each participant’s N1-peak topography, computed as the average absolute value per channel in the time interval from 0.14 to 0.18 s following the standard (see topography shown in Figure 4B). We then multiplied the time-domain data at all latencies and channels with these weights and averaged over channels, resulting in one virtual channel.

**Fig 3.**
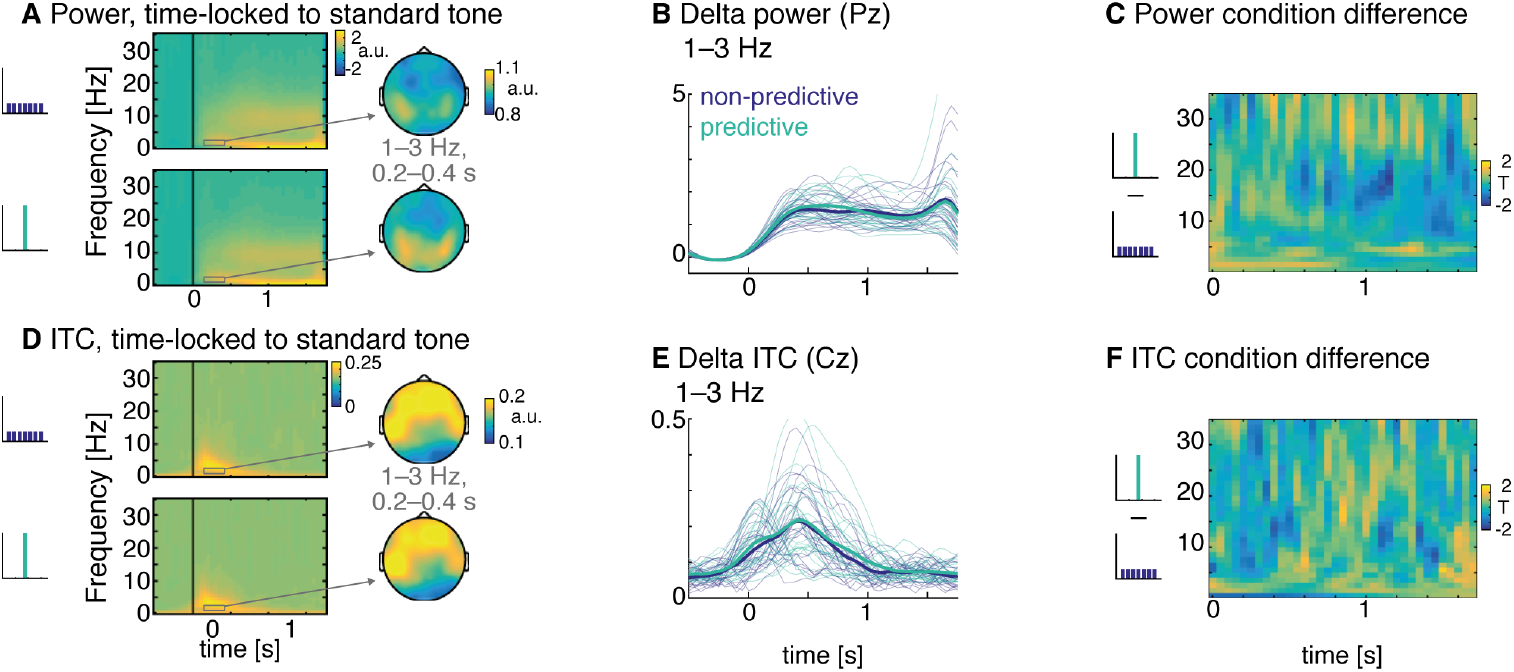
Time-frequency representations. **A. Power, time-locked to standard-onset**. Power estimates were baseline-corrected to the pre-standard interval and display relative change. Top panel: non-predictive condition, bottom panel: predictive condition. The topographies show the power scalp distributions in the interval from 0.2–0.4 s for frequencies from 1–3 Hz. **B. Delta power (1–3 Hz) over time** for the non-predictive (blue) and predictive conditions (green). Fine lines depict single participant s power values. **C. Power-difference between conditions (T-values)**. No significant condition differences were found. **D. Inter-trial phase coherence (ITC), time-locked to standard-onset**. Top panel: non-predictive condition, bottom panel: predictive condition. The topographies show the ITC scalp distributions in the interval from 0.2–0.4 s for frequencies from 1–3 Hz. **E. Delta ITC (1–3 Hz) over time** for the non-predictive (blue) and predictive conditions (green). Fine lines depict single participant’s ITC values. **F. ITC-difference between conditions (T-values)**. No significant condition differences were found.

**Fig 4.**
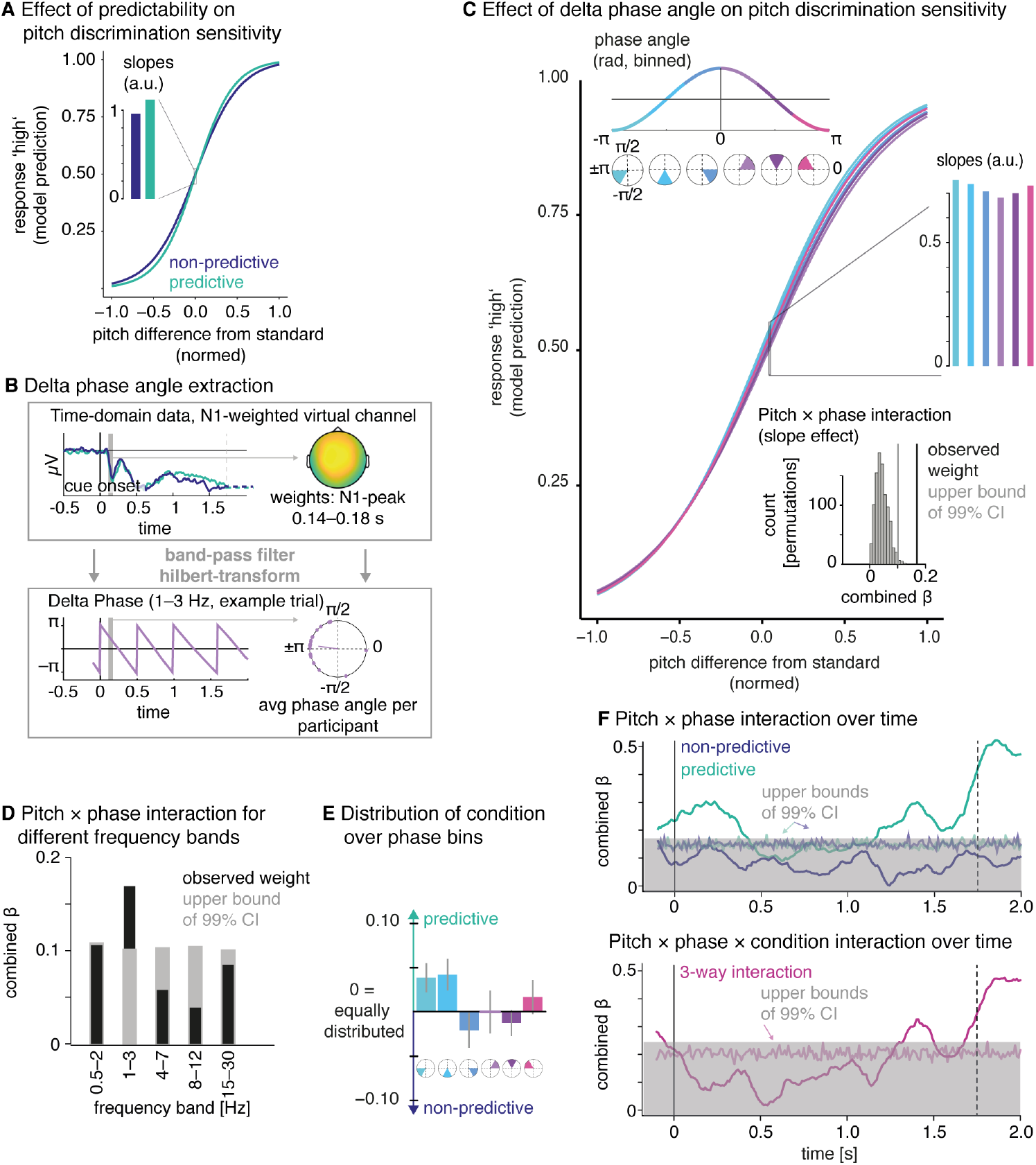
Delta phase angle predicts pitch discrimination sensitivity. **A. Replication of the behavioural effect** (s. Figure 1) with a logistic regression approach. Model predictions from the logistic regression with the predictors pitch (abscissa) and condition (colors). As illustrated by the bar-plot, there was a slope difference between conditions (i.e. an interaction between pitch and condition), with steeper slopes for the predictive condition. **B. Schematic depiction of the delta phase angle analysis**. We extracted the time domain data from single trials, from one virtual channel that reflects the weighted sum of the standard-evoked N1 topography (computed in the interval from 0.14–0.18 s), band-pass filtered (1–3 Hz) and applied the Hilbert transform, to extract the instantaneous phase angles in the time-window of 0.14–0.18 s (the N1-peak). **C. Effect of delta phase angle on pitch discrimination sensitivity**: Model predictions from the logistic regression model with the predictors pitch (abscissa) and phase angle (colors, binned only for visual display). There was a significant interaction between pitch and phase, that is the slopes of the psychometric functions differed depending on delta phase angle (depicted in the bar plot). Note that this analysis was performed on all trials, without separation into conditions. The inset on the bottom right side shows the observed interaction weight (in black) compared to a permutation distribution and its 99% confidence interval (in grey). **D. Pitch × phase interaction and confidence intervals for different frequency bands**. The grey bar shows the 99% confidence interval, the black bar the observed weight. Only for the delta band (1–3 Hz) the observed weight significantly exceeded the permuted weights. **E. Distribution of conditions over phase angles**. Conditions were coded as −1 for the non-predictive and 1 for the predictive condition, therefore an equal distribution of conditions over phase angle bins should result in an average condition (colored bars) of 0, which was not the case. Instead, more trials from the predictive condition occurred at the phase angles that were related to a steeper slope of the psychometric function (panel C). **F. Upper panel: Pitch × phase interaction over time, separated by condition**. The thick lines indicate the regression weights for the interaction over time for the predictive (green) and non-predictive condition (blue), the thin lines and grey shade indicate the 99% confidence interval computed with the permutation approach. **Lower panel: Condition × pitch × phase interaction over time**. The three-way interaction was significant only in the pre-target time window, indicating that only in the predictive condition delta phase angles predicted pitch discrimination performance during this time.

We applied a band-pass filter to the data (3rd order Butterworth, two-pass), with cut-off frequencies of 1 and 3 Hz for the delta band. After filtering, we applied the Hilbert transform and extracted phase angles as the imaginary value of the complex fourier spectrum averaged over latencies from 0.14 to 0.18 s, the peak latency of the N1. We chose the peak of the N1 as the window of interest, as the time point at which we measure the first reaction to the standard tone, possibly reflecting a phase reset of ongoing oscillations. Note that we did not choose the later time window in which the difference in the standard-evoked ERP significantly differed between conditions to avoid biasing our analysis for a between-condition effect.

We subjected the phase angles to a logistic regression to test for an effect of phase angle on the behavioural response, using the lme4 package in R (function glmer with a binomial link function, Bates et al., 2015). Per trial, we predicted the participant’s response in the pitch discrimination task (second tone lower or higher) with two numerical predictors, (1) the normalized pitch difference between standard and target tone (Δpitch in eq. 1, range −1-1, a.u.), and (2) the standard-evoked phase angle extracted as described above (*φ*), plus their interaction.

The predictors of the logistic regression can be interpreted following the logic of the psychometric function (DeCarlo, 1998), which models a behavioural measure (on the ordinate) based on variations of a stimulus feature (on the abscissa), and is described by two main parameters: threshold and slope. A threshold effect, that is a horizontal shift of the psychometric function, would be reflected by a main effect of the predictor *φ*, indicating a response bias, which we did not observe in the behavioral data. A slope effect, reflecting a shift in the steepness of the psychometric function, would result in an interaction between the predictors Δpitch and *φ*. Here, we were particularly interested in a slope effect, that is an interaction between the predictors pitch and phase angle. Due to computational constraints, we only specified a random intercept, but no random slopes for the predictors.

To account for the circularity of the phase angles, we followed an approach previously described by Wyart et al. (Wyart et al., 2012) (see also (Barne et al., 2017; Cravo et al., 2013)), using the sine and cosine of the phase angles jointly as linear predictors in a regression. For both, the *sin*(*φ*) and *cos*(*φ*), we specified an interaction with Δpitch:

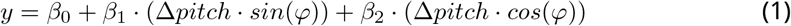

Then, we recombined the regression weights obtained for the interactions of *sin*(*φ*) and *cos*(*φ*) with Δpitch:

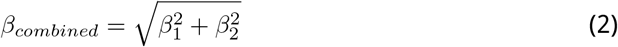

The resulting *β_combined_* is always positive and can thus not be tested against zero. We computed a reference distribution of *β_combined_* based on 1000 permutations, by permuting, per participant, the response values over trials, recomputed the model and retained the *β_combined_*. To assess significance of the interaction between pitch and phase angle, we assessed 99% one-sided confidence intervals, and computed p-values from the permutation distribution (Phipson and Smyth, 2010):

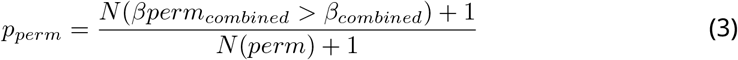

To visualize the modulation of pitch discrimination sensitivity over phase angles, we predicted responses from the logistic regression model (using the R package *emmeans,* Lenth, 2018), for a range of Δ*pitch, sin*(*φ*), and *cos*(*φ*) values, and plotted the resulting values for the recombined and binned *φ* (shown in Figure 4C).

We additionally computed the phase analysis on data filtered for the low delta (0.5–2 Hz), theta (4–7 Hz), alpha (8–12 Hz), and beta (15–30 Hz) frequency bands and tested the resulting *β_combined_* for significance using the permutation approach (Figure 4D). P-values were Bonferroni-corrected (accounting for five tests with a p-value threshold of 0.05, one for each frequency band), resulting in an adjusted alpha level of 0.01.

Furthermore, we assessed the time-course of the regression weights per condition by independently computing the model (Eq. 1) for each time point from −0.1 to 2 s and for each of the two conditions separately (Figure 4F). Here, we did not mute the time-domain data at target onset, since the model was computed separately per condition. To test for significance, we applied the permutation approach described above, using 200 permutations only (due to the time-consuming procedure). Finally, to test for condition differences, we computed the time-resolved logistic regression for both conditions jointly and added the factor condition to the above-described model to test for a three-way interaction.

### Distinguishing oscillatory from aperiodic activity

To assess whether the activity observed in the delta band is truly oscillatory, rather than reflecting aperiodic 1/f activity we applied irregular resampling (IRASA; Wen and Liu, 2016; see also Helfrich et al., 2018; Henry et al., 2016). This technique consists in downsampling the data at pairwise non-integer values and computing the geometric mean of the resulting power spectra. The resampling leaves the 1/f activity intact but removes narrow-band oscillatory activity. Once the 1/f activity has been obtained, it can be subtracted from the total power spectrum to assess only the oscillatory activity. We applied IRASA to the trial-wise data time-locked to the standard tone (–3 to 6 s), to the trial-averaged data per participant (ERP), and to 9 s of simulated data with a brown noise spectrum (see Figure 5A), as well as to single trial data from a 3 s snippet during the inter-trial interval (see Figure 5B). Power spectral density (PSD) was computed in sliding windows of 3 s in 0.25 s steps, using a fast Fourier transform tapered with a Hanning window for a frequency range of 0.33 – 25 Hz, without detrending, and the default resampling parameter (1.1 to 1.9, 0.05 increment). The PSD was normalized by dividing all values by the maximum value of the respective total PSD (trial data, ERP, and simulated data).

**Fig 5.**
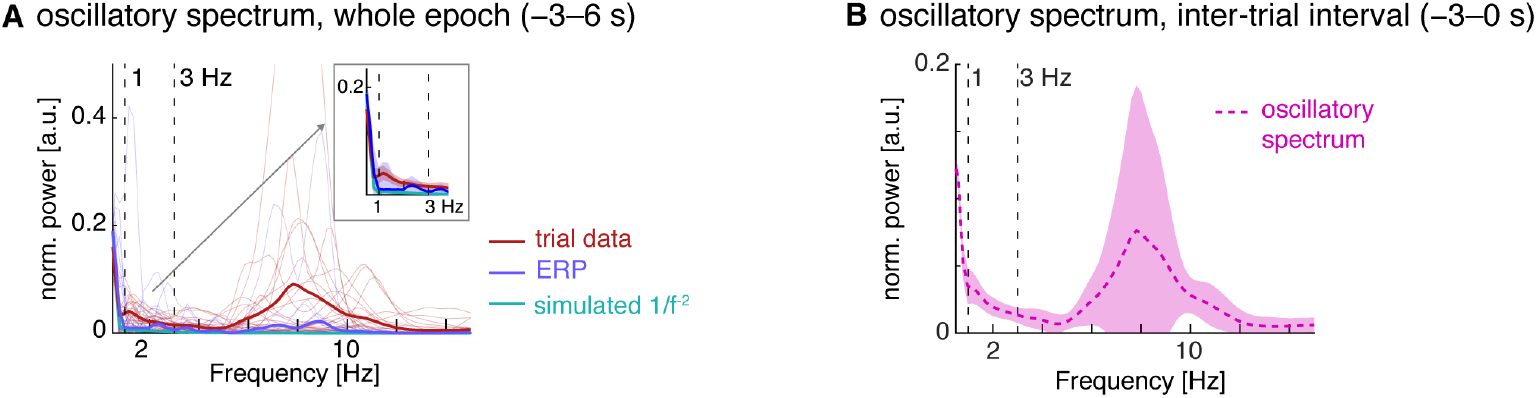
Testing for oscillatory activity in the 1–3 Hz range using the irregular resampling method. **A:** from single trial data (red), trial-averaged data (blue) and simulated brown noise (thick lines: average, fine lines: single participants). The left panel shows the oscillatory activity, obtained by subtracting the fractal PSD from the total PSD. The inset magnifies the delta frequency range from 1–3 Hz, and the shaded areas show 99% confidence intervals computed from a t-distribution. The difference between the red and blue lines shows that single trials contain additional, non-phase locked oscillatory activity in the 1–3 Hz band as compared to the ERP (trial average). **B:** Oscillatory spectrum obtained from resampling the pre-stimulus time window (3 s, taken from the ISI). Note that there is significant oscillatory activity in the 1–3 Hz range.

## Results

### Temporal predictability improves pitch discrimination

On average, participants’ responses were correct in 86% percent of trials. Using the full sample of 49 participants, we found that accuracy was significantly higher in the predictive compared to the non-predictive condition (T(48)=3.77, p<0.001, BF = 89.6); Figure 1B). We found a marginally significant increase in accuracy at the intermediate foreperiod for the predictive compared to the non-predictive condition (T(48)=1.8, p = 0.07, BF = 0.93); Figure 1B).

We furthermore analysed response times between conditions and over foreperiods. Response times were faster in the predictive (average 0.85 s), compared to the non-predictive condition (0.92 s), by about 70 ms (T(48)=8.3, p < 0.001, BF = 1^10^). As shown in Figure 1B, the difference is strongly driven by slower responses at shorter foreperiods in the non-predictive condition, but there was still a significant difference between the response times at the intermediate foreperiod only (T(48)=2.10, p = 0.04, BF = 1.47).

For the psychometric functions (depicted in Figure 1C), we observed a steeper slope in the predictive compared to the non-predictive condition (T(48)=3.85, p<0.001, Bayes Factor (BF)=114.3); Figure 1D), but no threshold effect (T(48)=1.05, p = 0.30, BF = 0.35); Figure 1E), nor effects on the lower asymptote (p = 0.48, BF = 0.27) or higher asymptote (p = 0.44, BF = 0.28).

To test whether the slope effect might be driven by shorter or longer foreperiods only, we computed psychometric functions on the trials with intermediate foreperiods (1.25–1.5 s in the behavioral sample, 1.33 – 2.17 s in the EEG sample; see Figure S1). We found a smaller but significant slope effect between conditions (T(48)= 2.73;p<0.01; BF = 5.46) showing that the slope difference was not solely driven by the shortest or longest foreperiods. Together with the condition differences in accuracy (not significant) and response times at the intermediate foreperiod only, this suggests that the performance improvement occurred not only at unexpectedly early or late foreperiods, but results from the difference in temporal predictability between conditions.

All of the above results held, albeit somewhat weaker, when analysing only data from participants for whom we had recorded EEG: Predictability resulted in marginally higher accuracy, (T(25)=1.82, p = 0.08, BF = 1.07), significantly larger PMF slopes (T(25)=2.60, p = 0.02, BF = 4.04), and no effects for the threshold, guess, and lapse rate (all p > 0.18, BF: 0.43, 0.61, 0.29, respectively).

To parallel the analysis of delta phase angles reported below, we also computed a logistic regression for the behavioural data, for the participants from the EEG sample only, with the predictors pitch difference (Δpitch), condition, and their interaction (plus random effects for all three). The analysis confirms the results described above, namely a significant main effect for Δpitch (p < 0.001), no main effect for condition (p = 0.9), but an interaction between Δpitch and condition (p < 0.01), which can be interpreted as a slope effect (see Figure 4A).

Finally, we assessed to what extend the predictability manipulation had been noticed by participants. During debriefing, no participant spontaneously reported to have noticed the manipulation of temporal predictability. Four participants from the behavioral and eight participants from the EEG sample said they had noticed the manipulation *after* the experimenter explained it. 16 (70%) of the behavioral and 17 (65%) of the EEG participants guessed correctly whether the high or low tones were temporally predictive in their case. Neither the participants who recognized the manipulation once it was explained, nor the ones who guessed correctly which tones were temporally predictive in their case showed a larger behavioral slope difference than the other ones (one-tailed Wilcoxon signed rank test, p = 0.88, p = 0.94, respectively). This suggests that the fact that participants were able to recognize the manipulation once it was explained did not reflect active engagement in timing during the experiment. The high percentage of correct guesses can possibly be explained by reverse inference, in which participants noticed that one condition was easier than the other, and – after learning about the predictive foreperiods – associated the perceived facilitation with predictability.

### Temporal predictability affects both, standard- and target-evoked event-related potentials

#### Standard-evoked activity

Event-related potentials were examined time-locked to the standard-tone (Figure 2A). Both conditions showed a negative deflection between 0.1–0.2 s after the standard onset, with a peak at 0.16 s and a fronto-central topography. We refer to this component as the standard-evoked N1. We observed a significant difference between conditions in the time window of the late N1/ early P2 component, where amplitude was more negative for standards that were temporally predictive of the onset of the target (predictive condition; 0.21–0.26 s, p = 0.02). This difference is important in that it shows that standard tones were processed differently if they served as a temporal cue for the target onset versus did not serve as a temporal cue. The latency and topography of the standard-evoked N1 (not the time-range in which the difference was found which was slightly later) was used for the analysis of phase angles described below. When directly comparing the ERPs evoked by the 550 versus 950 Hz standards (randomly assigned to the predictive and non-predictive condition over participants), there was no statistically significant difference in the early time window following the standard tone.

#### Target-evoked activity

Event-related potentials time-locked to the target-tone (Figure 2B) also showed a negative deflection between 0.1–0.2 s after the target onset, with a fronto-central topography. We refer to this component as the target-evoked N1. For targets in the predictive condition, the N1 was larger (0.09–0.14 s, p = 0.02). Importantly, the difference is not solely due to the onset time of the target (see inset in Figure 2B and Figure S2), which would be reflected by a difference only for long or short foreperiods in the non-predictive condition.

To test for an apparent latency shift in the N1 between the non-predictive and predictive conditions, we computed the half-area measurement (Luck, 2005), which indexes the time-point at which half the area of a deflection has been reached. Compared to peak-latencies, this measure accounts better for asymmetric deflections. We found a significantly earlier N1-latency for the predictive, compared to the non-predictive condition (Cz, 0.13 s versus 0.15 s; T(23)=3.03, p < 0.01).

Furthermore, there was an amplitude difference at a later positive prolonged component, which was positive at posterior and negative at frontal electrodes (0.20–0.45 s, p<0.01; 0.28–0.36 s, p = 0.02). When computing the analysis using only trials with foreperiods ≥1.75 s (and equating the number of trials in the predictive condition for a fair comparison), the early cluster and the later frontal clusters remained (0.09–0.14 s, p = 0.04; 0.25–0.37 s, p = 0.008, marked in light blue in Figure 2B, right panel). When running the same analysis on the trials ≤1.75 s, we again found the early cluster (0.08–0.14 s, p = 0.01), and the later posterior cluster (0.16–0.49 s, p<0.001, marked in pink in Figure 2B). These findings show that the early difference was not driven by the shorter or longer foreperiods separately, but resulted from temporal predictability per se. The positive difference at posterior channels (cluster marked in pink in Figure 2B), however, was driven by the short foreperiod trials, and the negative difference at frontal channels (cluster marked in light blue in Figure 2B) was driven by the long foreperiod trials.

### No condition differences in delta (1–3 Hz) power or ITC during the foreperiod

We assessed power in a frequency range between 0.5–34.5 Hz for the predictive and non-predictive conditions (see Figure 3A), time-locked to standard onset. Both conditions showed an increase in power in the delta-range (1–3 Hz, Figure 3B) after standard onset, and a prolonged increase in the alpha-range (8–12 Hz) relative to baseline. We found no statistically significant power differences between conditions at the cluster level (see Figure 3C).

ITC across the 1–3 Hz range did show the expected increase following the standard tone, ranging from 1–3 Hz, and a prolonged increase in the delta band in both conditions (Figure 3D,E). However, when comparing inter-trial phase coherence (ITC) for all frequencies between conditions, no significant differences were observed. A hypothesis-driven cluster test restricted to the delta frequency band (1–3 Hz) revealed a non-significant cluster of enhanced delta ITC (Figure S3; 0.85–1.1 s, 1.5–2.5 Hz, p = 0.19). This shows that delta ITC increased nominally, albeit not significantly in the predictive condition. Likely the effect is too weak to reach significance either because of signal processing constraints (muting of target-evoked activity), or the absence of an entraining rhythm.

### Standard-evoked delta phase angle predicts pitch discrimination sensitivity

To test whether delta oscillations play a role in temporally predictive processing in this study, we tested for a relation between delta phase angles and pitch discrimination performance using a logistic regression approach (see Figure 4B for a schematic depiction). A timing mechanism that predicts the onset of the target tone would have to be initiated at the standard tone, which acts as a temporal cue, which is why we were particularly interested in this time window. We chose the peak of the N1 as time point of interest, as it is the earliest measurable response to the temporal cue. We hypothesized that temporal predictions could possibly be implemented via a phase reset of an ongoing delta oscillation.

Phase angles in the post-standard time window (0.14–0.18 s) were extracted by applying the Hilbert transform to band-pass filtered (1–3 Hz) single trial data with one virtual channel (see Methods for details) representing the sum of all channels weighted by the N1-topography. We subjected the phase angles (as their sine and cosine) to a logistic regression with two numerical predictors, the normalized pitch difference between standard and target tone, and the standard-evoked phase angle, plus their interaction. To assess significance of the interaction effect, we used a permutation approach. We found a significant interaction between pitch and phase angle, which indicates that the slope of the psychometric function varied depending on the delta phase angle evoked by the standard tone (Figure 4 C). The interaction effect was significant only for the delta band (1–3 Hz), but not for other frequency bands tested (0.5–2 Hz; 4–7 Hz; 8–12 Hz; 15–30 Hz; Figure 4 D). Note that this procedure was performed on all trials, without separation into conditions, and thus is generally valid, both for trials on which the standard served as a temporal cue and trials for which it did not.

Next, we tested whether the interaction between delta phase angle and pitch discrimination sensitivity was specifically driven by our manipulation of temporal predictability. We examined the regression weight for the interaction at different time points over the trial, and independently for the predictive and non-predictive conditions. This analysis (Figure 4F, upper panel) showed that the interaction effect between delta phase angle and the slope of the psychometric function was significant (i.e. exceeded the 99% confidence interval of the permutation distribution) only for the predictive condition, and occurred at two time points: after the standard tone (around 0–0.4 s), and prior to target onset (around 1.1–1.4 s). We therefore conclude that the interaction effect was mainly driven by the predictive condition.

The three-way interaction between condition, delta phase angle, and pitch discrimination sensitivity was significant only in the later time window (Figure 4F, lower panel). A supplementary analysis testing the effect of different foreperiods (target onset times) on delta phase angles in the non-predictive condition (Figure S4), confirmed that phase angles in the time ranges in which we observed the above-described effects were not affected by the different target offsets in the non-predictive condition.

We also assessed the relationship between phase angle (binned into 6 bins for this purpose) and condition (indexed as −1 for the non-predictive and 1 for the predictive condition; Figure 4 E). If the trials would be equally distributed over conditions per phase angle bin, this should result in an average condition of 0 at all phase angles. We found more trials of the predictive condition to occur at the phase angles at which we had found the higher slopes (Figure 4 C), which suggests that phase angles varied between the two conditions. A post-hoc test for a quadratic effect of phase bin on condition (computing a generalized linear model predicting condition from phase bins) yielded only a marginally significant weight for this contrast (p = 0.09). We thus conclude that there is no meaningful phase angle difference between conditions at the population level.

### Additional analyses

#### Oscillatory versus 1/f activity

To test for the presence of oscillatory activity in the delta band, we subtracted fractal power spectra (obtained using the irregular resampling method (IRASA; Wen and Liu, 2016) from the total power spectra. The results (depicted in Figure 5, see also S5) show that albeit no clear peaks can be found in the delta range, power spectral density (PSD) computed from single trial data was higher in the 1–3 Hz range compared PSD computed on the ERP and simulated data (Figure 5A). If anything, the PSD computed on single trial data has a small peak around 1 Hz, while the PSD of the ERP has two smaller peaks at 3 and 4 Hz. When computing the same analysis on pre-stimulus data (from the ISI, 3 s signals), we observe residual oscillatory activity in the 1–3 Hz range (Figure 5B). While it is difficult to completely separate oscillatory from 1/f activity at slow frequencies - and to our knowledge, no previous study showed a clear oscillatory peak in the PSD in the delta range – our analyses suggest some oscillatory activity in the delta band.

#### Mediation analysis

We also considered mathematically the possibility that delta phase angle in the post-cue time window would mediate the effect of temporal predictability on pitch discrimination sensitivity, by comparing the regression weight of the interaction between pitch and temporal predictability estimated from a model with no other predictors (as depicted in 4A), and from a model that additionally contained an interaction term for pitch and phase angle (Baron and Kenny, 1986; Muller et al., 2005). The negligible change in weight between both models (0.307 to 0.304) indicates that there is no evidence for a mediation effect.

#### Delta phase versus ERP effect

To distinguish between the ERP effect (found on the N1) and the delta phase effect, we tested whether the N1 amplitude could explain the findings. Computing the same logistic regression model with the N1 amplitude (averaged activity between 0.14–0.18 s, using the same spatial filter) instead of the phase angles as above revealed no significant interaction effect (p = 0.15), i.e. the N1 amplitude does not predict pitch discrimination performance on single trials and can thus not simply replace the delta phase angle. However, the N1 amplitude correlated significantly with the standard-evoked phase-angle at all frequency bands, as assessed by a circular-linear correlation (from the Directional package in R, Tsagris et al., 2018); *R*^2^: 0.5–2 Hz: 0.21, 1–3 Hz: 0.27, 4–7 Hz: 0.06, 8–12 Hz: 0.056, 15–30 Hz: 0.004 (all p-values <0.001).

## Discussion

In this study, we asked whether human listeners extract implicit temporal contingencies from auditory input to form temporal predictions in absence of a periodic input structure. If so, how are such endogenous temporal predictions represented in neural dynamics? We implicitly manipulated temporal predictability by varying the foreperiod (i.e., the interval between standard and target tones) in a pitch discrimination task. Unbeknownst to participants, one of two possible pitches used as the standard tone was indicative to one of two foreperiod distributions, respectively: drawn either from a uniform distribution, under which the onset of the target tone was unpredictable, or from a single-valued distribution under which the onset of the target tone was fully predictable.

The data reveal several indices that participants did form temporal predictions: most importantly an increase in pitch discrimination sensitivity in the predictive condition, and condition differences in the evoked response to standard- and target tones. However, contrary to our initial hypothesis, classical time-frequency analyses revealed no differences in power or inter-trial phase coherence in slow oscillatory frequencies. Yet, a direct analysis of delta phase angles shows that the phase of delta oscillations in response to the standard tone and in the pre-target time window is indicative of pitch discrimination performance. This finding suggests an instrumental role of delta oscillations in implementing endogenous temporal predictions for audition.

### Implicit temporal predictability improves pitch discrimination sensitivity

Behaviourally, we observed an increase in pitch discrimination sensitivity in the temporally predictive condition, reflected in a steeper slope of the psychometric function (Figure 1). Even though the absolute difference in behaviour is not large, we observed a robust set of converging effects of temporal predictability on response times, accuracy and slopes (49 participants). These suggest that listeners can implicitly learn to associate sensory stimulus features like pitch with single-interval temporal predictions, emphasizing the relevance and ubiquitousness of timing in human cognitive processing.

Importantly, participants were not made aware of the predictability manipulation, and no participant was able to correctly describe it during debriefing. About 25% of participants were able to recognize the manipulation after it was described by the experimenter, but did not show a larger behavioural effect, suggesting they did not actively engage in timing. The fact that a majority of participants guessed correctly which standard tone was associated with temporal predictability can be explained by reverse inference: participants noticed that one condition was easier than the other, and - after being informed about the predictive foreperiods during debriefing - associated the perceived facilitation with predictability.

To our knowledge, this is the first study to show that pitch discrimination sensitivity is improved by implicit but non-rhythmic temporal predictions. In the auditory domain, detection speed and performance are facilitated by rhythmic temporal predictability (Henry et al., 2014; Henry and Obleser, 2012; Lawrance et al., 2014; Wright and Fitzgerald, 2004), but the use of detection tasks might underline the timing aspects of the task.

One previous study (citepbausenhart2007knowing showed that shorter presentation times (difference of about 6 ms) are needed for to achieve correct pitch discrimination performance, when the target tone occurs with a block of constantly short foreperiods. Another study (Morillon et al., 2016) revealed that aperiodic regularities improved auditory sensitivity when participants had to discriminate a deviant tone from standards, but likely the manipulation was more easily detectable by participants due to the use of rhythmic and monotonically increasing intervals. Complementing these previous findings, we here show that implicit temporal predictability improves auditory perceptual processing in absence of an embedding rhythm, and despite any explicit incentive to engage in timing.

### Temporal predictions affect neural processing of predictive and predicted tones

#### Predictive tones (standards)

An important indicator for the successful extraction of temporal predictability is the difference in event-related potentials evoked by predictive and non-predictive standard tones (Figure 2A). It suggests that participants learned to associate the pitch of the standard tone to temporal predictability, and flexibly used the standard as a temporal cue on a trial-by-trial basis.

Few studies have investigated effects of predictability on the early sensory processing of the *predictive* or cue stimulus. In spatial cueing, there is evidence for an effect of predictions on early positive and negative cue-evoked components (100–200 ms post cue; Jongen et al., 2007; Nobre et al., 2000; Yamaguchi et al., 1994). In the temporal domain, there is, to our knowledge, only one study that showed an enhanced N1-after temporal cues (in 8-12 years old children, Mento and Vallesi, 2016). Our results are in line with this finding and reveal that the cue-evoked N1 in adults is affected even by implicit temporal predictability. The observed N1 enhancement, a response previously assigned to the recruitment of additional attentional resources in the context of predictive processing (Bendixen et al., 2012), could speculatively be explained as enhanced attentional processing of the predictive cue, which conveys more information about future task-relevant events than a non-predictive cues.

#### Predicted tones (targets)

In response to target tones, we found a larger and faster N1 in the predictive compared to the non-predictive condition, suggesting a facilitation of sensory processing of temporally predicted targets (Figure 2B). This result corroborates a large base of studies reporting mainly amplitude effects of temporal predictability on sensory evoked potentials (Correa et al., 2006;Hsu et al., 2014;Hughes et al., 2013; Kok et al., 2011; Lampar and Lange, 2011; Lange, 2009; Miniussi et al., 1999; Sanders and Astheimer, 2008; Schwartze et al., 2013). The reported direction of those amplitude effects varies with the paradigm used (for an extensive discussion see Lange, 2013) - for probabilistic foreperiod variations as used here, both, reduced (Paris et al., 2016; Sherwell et al., 2017) and enhanced N1 amplitudes (Griffin et al., 2002) have been reported.

The main specificity of the present study is that we only manipulated temporal, not spectral predictions, and hence the temporal prediction could have resulted in a faster and more efficient allocation of attentional resources to predicted stimuli, to facilitate the assessment of their pitch, which could not be predicted.

The observed latency-shift of the N1 by temporal predictions is in line with one previous study using a manipulation of foreperiods (Seibold et al., 2011), and one study on rhythmic temporal predictability (Rimmele et al., 2011). Further evidence comes from experiments reporting a faster N1 for auditory speech and non-speech events combined with visual events (Paris et al., 2017; Stekelenburg and Vroomen, 2007; Vroomen and Stekelenburg, 2010; Wassenhove et al., 2005). Note that in our study, the predictive information conveyed by the cue was purely temporal, since the pitch of the target tones was unpredictable. In sum, the facilitation of the target-evoked N1 suggests that temporal predictions alone can enhance early auditory processing.

### Implementation of temporal prediction through slow neural oscillations

A central aim of this study was to assess the role of slow neural oscillations for an endogenous representation of temporal predictions. Previous studies convincingly established a mechanism of facilitation of sensory processing via phase alignment of delta oscillations for stimuli that occur during the preferred phase, i.e. in synchrony with the preceding rhythm (Cravo et al., 2013; Henry et al., 2014; Kösem et al., 2018; Lakatos et al., 2008; Schroeder and Lakatos, 2009). An open question is however, whether the alignment of slow neural oscillations towards predicted stimulus onsets is contingent on rhythmic entrainment to the exogenous stimulation, or whether slow oscillations also implement endogenous temporal predictions, for example via single-trial phase resets.

We found no robust condition differences in oscillatory power or phase using classical time-frequency analyses (see Figure 3). The absence of condition differences in phase coherence during the foreperiod (Figure 3F) replicates our previous results (Herbst and Obleser, 2017) and suggests that enhanced phase coherence (Breska and Deouell, 2017; Cravo et al., 2011) might be affected by dedicated or residual periodicity in the stimulation (Obleser et al., 2017), and/or overt engagement in timing (Stefanics et al., 2010). As a side note, it is important to emphasize the methodological challenge of analysing low frequency oscillations in the pre-target window. The probabilistic manipulation of foreperiods as applied here results in differential time-locking of target activity between conditions, and our conservative approach of removing this activity might have weakened existing pre-target differences through back-smearing of the muted activity. Here, a nominal increase in delta phase coherence was found in the predictive condition (Figure S3), but failed to pass the threshold for statistical significance, suggesting that a phase coherence effect is not fully absent in non-rhythmic temporal predictions, but not strong enough to be measured with the available techniques. Thus, the representation of temporal predictions by enhanced phase coherence - or at least our ability to measure it in human EEG - is likely contingent on rhythmic stimulation.

Crucially, we found that the absolute phase angle of the delta oscillation in auditory areas shortly after the temporal cue predicted behavioural sensitivity in response to the later-occurring target tone (see Figure 4C). The effect was observed for data spatially filtered with a topography relevant for auditory stimulus processing (from the N1), suggesting auditory cortex as the most likely generator. Furthermore, the effect was specific for the delta band (1–3 Hz) with the highest sensitivity occurring at phase angles closest to the trough of the delta oscillation (±*π*) at the cue and about 1.4 s post-cue (average period of 0.5 s). Albeit interpreting the absolute phase angle from EEG data demands caution, this corroborates the idea that the trough of the delta oscillation is a particularly beneficial state for auditory perception (Henry et al., 2016; Lakatos et al., 2013). Theoretically, the proposed mechanism should surface as a relationship between delta phase and behavior throughout the whole foreperiod interval, but here we only observed it in the post-cue and pre-target time intervals. It is conceivable that the discontinuity of the effect throughout the foreperiod results from the EEG signal reflecting summed activity from large populations of neurons: other neural processes might overlay the maintenance of delta phase throughout the foreperiod, which surfaces only during the initiation of the prediction by the temporal cue, and the anticipation of the target, which are the most relevant time points for the proposed mechanism.

This relationship between delta phase and behavioural sensitivity held across all trials, regardless of their experimental condition. However, a follow-up analysis per condition found this relationship between delta phase angle in the post-cue time window and behavioural sensitivity to occur only in the predictive condition (Figure 4F, upper panel). To test whether the relationship between delta phase and behavioural sensitivity differed statistically between conditions, we computed the three-way interaction between pitch, delta phase angle, and condition (4F, lower panel), which was significant only during the pre-target time window. Possibly, low statistical power for this particular analysis prevented us from confirming the condition difference in the post-cue time window. This finding thus suggests that delta phase in the post-cue time window affects behavioural sensitivity in both conditions, while the effect found in the pre-target time window is specific to the predictive condition only.

This per se is not proof of a *causal* chain from temporal predictability *via* optimized phase angle of delta oscillations to increased auditory sensitivity. While not state of the art in neuroscience, our analysis did fail to establish hard statistical evidence for such a mediation effect. Possibly, different steps necessary to accommodate the complexity of our data in the model (dealing with the circular measure of phase angle and assessing an interaction effect as a measure of behavioural sensitivity), and the small proportion of variance explained by the experimental manipulation (a common problem in cognitive neuroscience) might have prevented us from observing a mediation effect (but see Benwell et al., 2017, for a successful example).

An important question is to what respect the observed phase effect reflects truly oscillatory activity, rather than a modulation of the evoked response to the standard or target tones. Admittedly, temporal smearing occurs due to the long analysis windows needed to capture slow oscillations. Importantly, the contingency between delta phase angle and auditory sensitivity re-occurs in the pre-target time window at around 1.4 s and does not rise monotonically into the post-target window. Therefore, we deem it unlikely this effect resulted from back-smearing of target-evoked activity.

Furthermore, the observed phase effect is specific to the frequency range identified by previous studies, rather than resulting from broad-band activity - as one would have expected from a purely evoked effect. We further showed that the N1 amplitude itself does not show the critical relationship with behavioural sensitivity, although the two measures correlate, arguing for a more specific role of delta oscillations in temporal prediction. In fact, the ERP might at least partially result from a reset of ongoing neural dynamics by the onset of a stimulus (Makeig et al., 2002).

The effect is strongest in the 1–3 Hz range, and not at the frequencies that would reflect the stimulation (0.57 Hz for the intermediate foreperiod of 1.75 s), which is in line with a study that showed selective entrainment at 1.33 Hz despite stimulation at 0.67 Hz (Gomez-Ramirez et al., 2011). These findings align with the assumption that auditory processing fluctuates with the phase of endogenous delta oscillations in the absence of evoked activity (Henry et al., 2016; Kayser, 2019; Kayser et al., 2015; Stefanics et al., 2010).

Not least, additional spectral analyses suggest some oscillatory activity in the delta band after subtracting the 1/f spectrum, which is not explained by the ERP (see Figure 5 and S5 for comparison of the spectra).

Taken together, these findings speak for a dedicated mechanism that implements temporal predictability in the auditory domain via a phase shift of auditory-cortical delta oscillations. While this study was not designed to directly test assumptions derived from dynamic attending theory (Jones, 1976;Large and Jones, 1999), but rather to assess the endogenous implementation of temporal predictions through a neural phase code, our findings are consistent with a dynamic adjustment of attentional windows to events in time.

We acknowledge that as an alternative explanation to an oscillatory effect, it is conceivable that the activity we observe reflects the extraction of temporal predictions from the temporal cue, but that another process is responsible for maintaining this prediction throughout the foreperiod interval to alert the system when it it is time to expect the target stimulus. For instance, this could be achieved via top-down projections from auditory areas towards thalamic and thalamostriatal pathways described as crucial for auditory timing (Barczak et al., 2018; Ponvert and Jaramillo, 2018), converging with an instrumental role of the striatum in explicit timing (Mello et al., 2015). Future research is needed to assess sub-cortical circuits.

In sum, our findings do underline the relevance and specificity of delta oscillations for an endogenous representation of temporal predictions. The adjustment of phase angles at the cue can be seen as the initiation of a timing process, which prepares the system to be in a beneficial state at an anticipated time point, resulting in an optimized delta phase angle prior to target onset.

## Conclusions

Human listeners do use strictly implicit temporal contingencies to perform a sensory task for which timing is not an explicit requirement. Here, we assessed how temporal predictions are implemented in neural dynamics by combining psychophysics and EEG data. We found endogenous temporal predictions for audition to be reflected in the phase of delta oscillations, likely via an optimized phase reset of delta oscillations in auditory areas evoked by a temporal cue. These results point towards an instrumental role of delta oscillations in initiating temporal predictions, even in the absence of an entraining rhythm.

## Supporting information

**Fig. S1.**
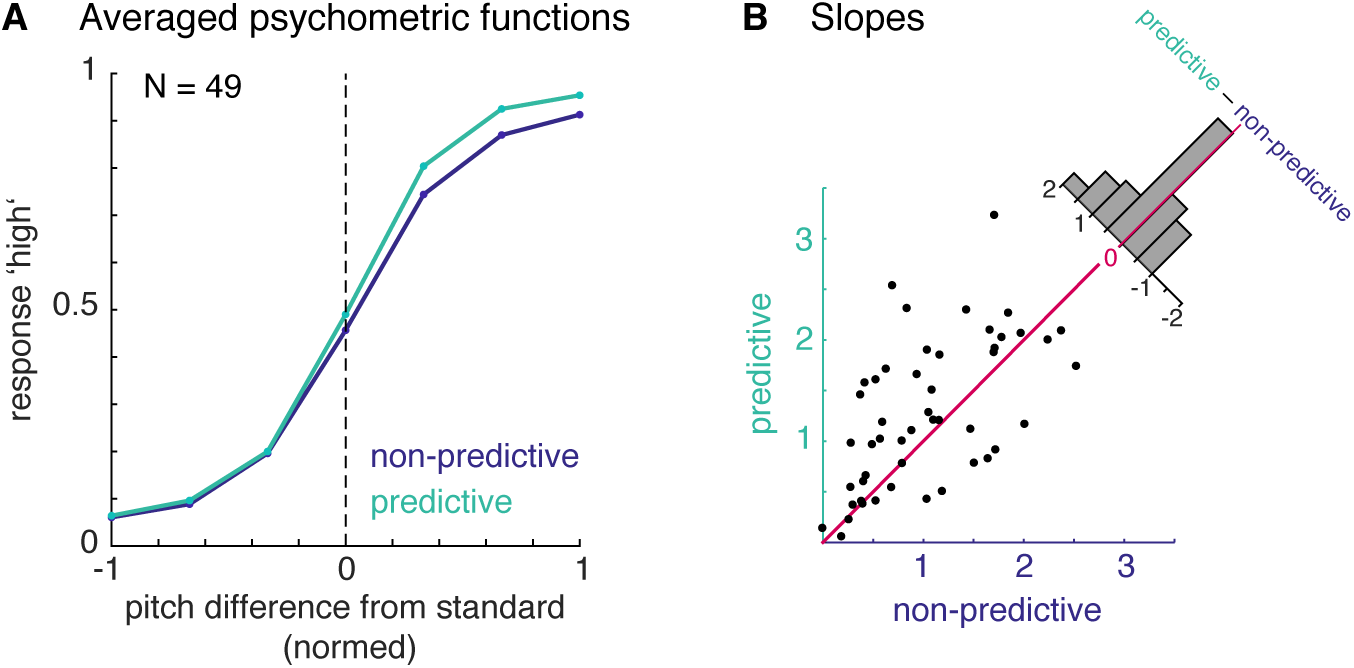
Slope effect at intermediate foreperiods. **A.** Psychometric curves, fitted only at a small range of intermediate foreperiods. **B.** Slopes for the predictive and non-predictive conditions at intermediate foreperiods only. This additional analysis was performed to rule out the possibility that the slope effect was solely driven by the shortest and longest foreperiods in the non-predictive condition.

**Fig. S2.**
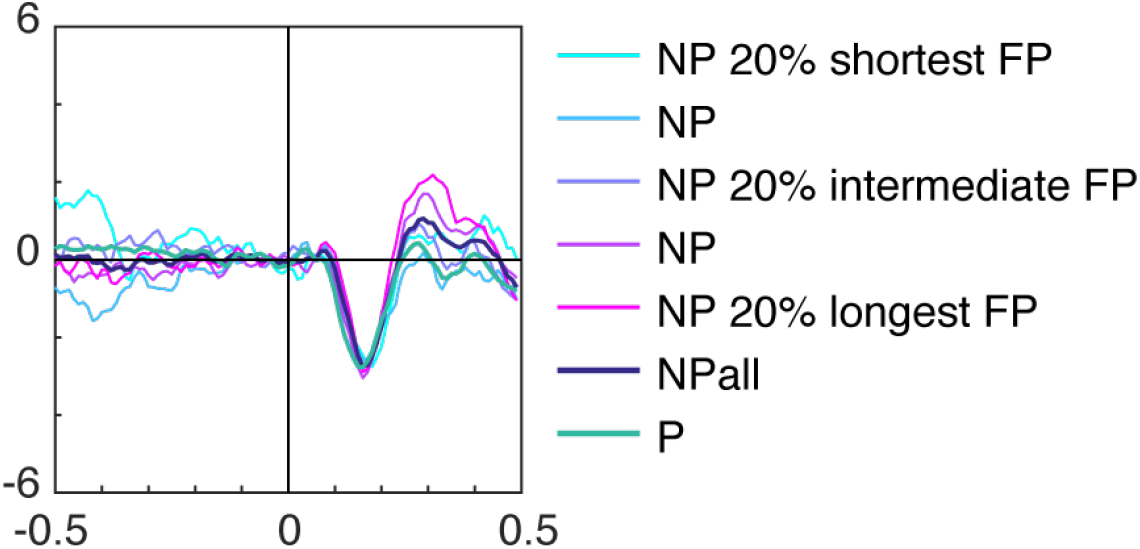
Target-evoked ERP by foreperiod. Target-evoked ERPs for the predictive (green) and non-predictive (dark blue) condition. The trials for the non-predictive condition were split into five foreperiod bins from the 20% shortest to the 20% longest foreperiods (cyan to pink).

**Fig. S3.**
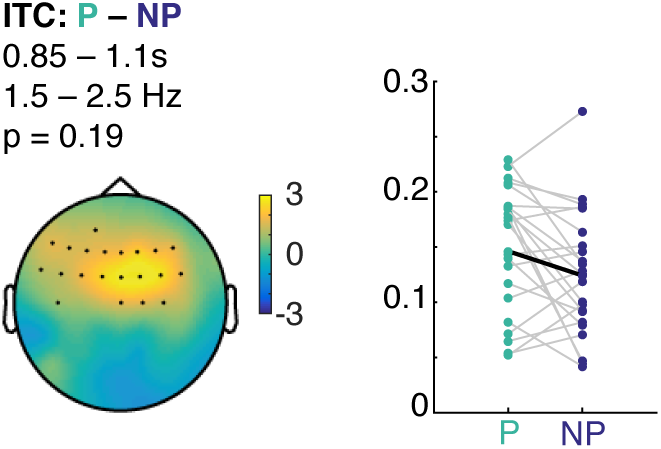
Hypothesis-driven cluster-test for a condition difference in delta ITC. We did not observe any statistically significant differences in delta ITC during the foreperiod, but a hypothesis-driven test restricted to the delta band showed a cluster that failed to pass the threshold for significance. This shows that there was nominally, albeit not significantly increased delta ITC in the predictive condition, but likely the effect is too weak either because of signal processing constraints, or its contingency on an entraining rhythm.

**Fig. S4.**
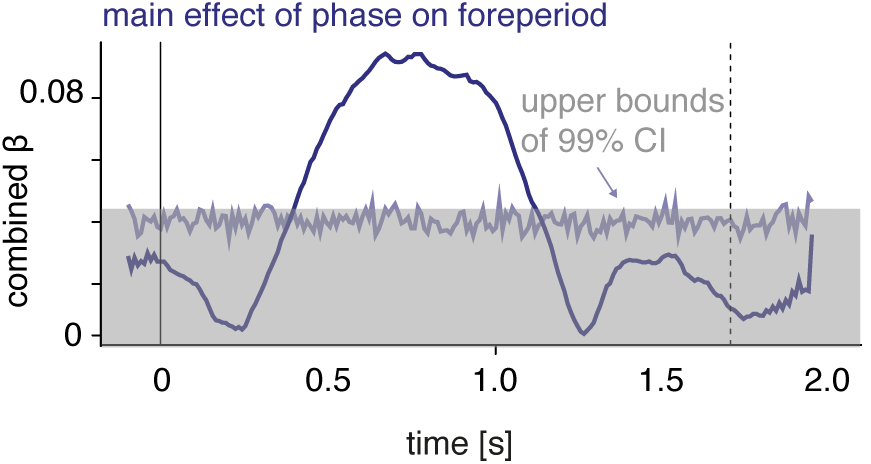
Predicting the foreperiod from phase angles. We tested whether delta phase angle time series in the non-predictive condition are affected by the different target onset times (i.e. foreperiods). To this end, we computed a linear mixed effect model, predicting foreperiod by phase angles, separately at each time point. Phase angles were separated into their sine and cosine and the *β_combined_* was tested against a permutation distribution (200 samples) for which the assignment between foreperiod and phase angles was randomized. The result shows a relation between phase angles and foreperiods in the time window between 0.5-1 s, but not in the time windows in which the critical effects depicted in Figure 4F were found.

**Fig. S5.**
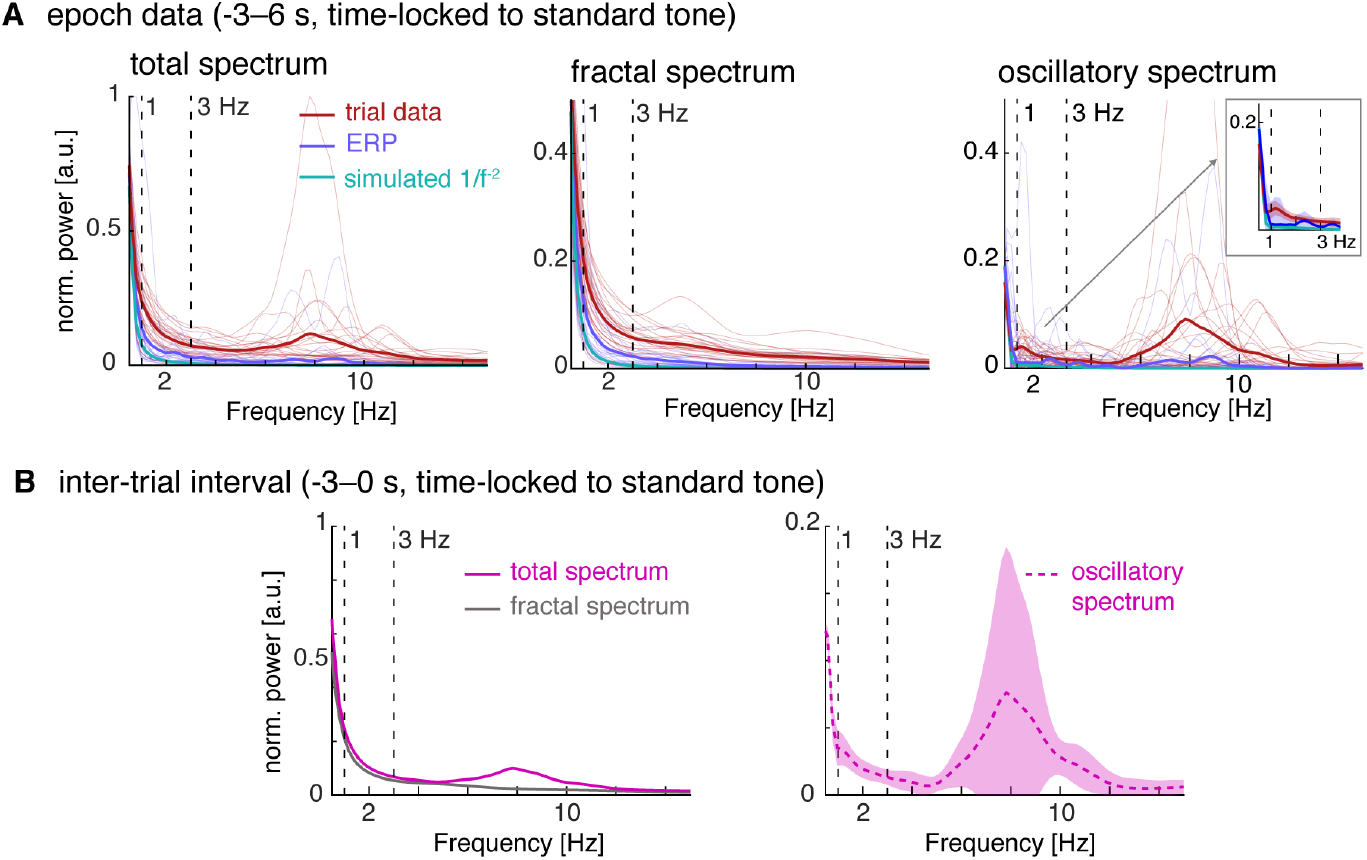
Power spectral density (PSD) computed using the irregular resampling method (IRASA; Wen & Liu, 2016). **A:** PSD of single trial data (red), trial-averaged ERP data (blue) and simulated brown noise (green;thick lines: average, fine lines: single participants). PSD were normalized by dividing all values by the maximum value of the respective total PSD (trial data, ERP, and simulated data). Nine second data snippets were used, time-locked to the standard tone. PSD was computed in sliding windows of 3 s in 0.25 s steps, using fast a Fourier transform tapered with a Hanning window for a frequency range of 0.33 – 25 Hz, without detrending. The left panel shows the total spectrum, computed as the auto-power spectrum of the respective input data. The middle panel shows the fractal spectrum, computed as the geometric mean of the auto-spectra of the pairwise resampled time-series (using the default resampling parameter: 1.1 to 1.9 with a 0.05 increment). The right panel shows the oscillatory activity, obtained by subtracting the resampled PSD from the total PSD. The inset magnifies the delta frequency range from 1–3 Hz, and the shaded areas show 99% confidence intervals computed from a t-distribution. **B**: Irregular resampling computed in a inter-trial interval (3 s). Left: total (pink) and fractal (grey) spectra;right: oscillatory spectrum with 99% confidence interval.

## Acknowledgments

This research was supported by a DFG grant (HE 7520/1-1) to SKH. The authors would like to thank Anne Herrmann for overseeing the data acquisition, Michael Ploechl for technical support, and Virginie van Wassenhove and the Cognition & Brain Dynamics Team at NeuroSpin for helpful discussions.

